# A Comparison of Drug Delivery into Skin Using Topical Solutions, Needle Injections and Jet Injections

**DOI:** 10.1101/694943

**Authors:** Katharina Cu, Ruchi Bansal, Samir Mitragotri, David Fernandez Rivas

## Abstract

Drug diffusion within the skin with a needle-free micro-jet injection (NFI) device was compared with two well-established delivery methods: topical application and solid needle injection. A permanent make-up (PMU) machine, normally used for dermal pigmentation, was utilized as a solid needle injection method. For NFIs a continuous wave (CW) laser diode was used to create a bubble inside a microfluidic device containing a light absorbing solution. Each method delivered two different solutions into *ex-vivo* porcine skin. The first solution consisted of a red dye (direct red 81) and rhodamine B in water. The second solution was direct red 81 and rhodamine B in water and glycerol. For PMU experiments, the skin samples were kept stationary and the diffusion depth, width and surface area were measured. The NFI has a higher vertical dispersion velocity of 3 × 10^5^ *μm*/s compared to topical (0.1 μm/s) and needle injection (53 *μm*/s). The limitations and advantages of each method are discussed, and we conclude that the micro-jet injector represents a fast and minimally invasive injection method, while the solid needle injector causes notably tissue damage. In contrast, the topical method had the slowest diffusion rate but causes no visible damage to the skin.

## 1. Introduction

For many centuries, needles and syringes have been extensively used in several medical procedures. And for just as long, injections are feared by many patients.^1, 31, 32^ Pills or topical skin products are easier to use and are painless. Accordingly, oral and transdermal administration routes are favored by patients and physicians alike.^3, 10, 18, 30^ However, injections are difficult to replace since certain drugs can only be administered via intramuscular, subcutaneous or intravenous injections, where drugs reach the systemic circulation with high efficiency. Topically applied drugs via creams or patches exhibit slow drug uptake due to the passive delivery across the skin induced by a concentration gradient, in which the diffusion properties are a function of the skin characteristics and the solution molecules.^32^ The slow diffusion originates primarily from the properties of the outermost skin layer, the stratum corneum (SC), which protects the underlying tissue from infections and dehydration. In practice, diffusion is only limited to lipophilic and low molecular weight drugs (< 500 *g*/*mol*).^4, 10, 32^ Thus, the SC poses a great permeation challenge for most of the drugs and delivery methods since the majority has a high molecular weight and poor solubility.^6^ One alternative to the topical application is the use of microneedles, which are effective for breaking through the SC but have poor accuracy of delivery, among other limitations.^20^

Other injection alternatives developed over the last decades, such as high-pressure injections and needle-assisted jet injectors, have shown advantages of needle-free injections (NFIs), in which a pressurized liquid (or powder) jet penetrates the skin.^5, 19, 22, 25, 26, 40^ Infections as well as disease transmissions due to improper (re)use of needles can become irrelevant when the disposal of needles is not required, which also reduces high costs for single-use components.^4, 12^ The most important advantage of an NFI device is thought to be pain reduction that results in higher patient compliance, especially in chronic diseases like diabetes or by people fearing sharps.^16, 17^ From the different energy sources used to power NFIs, a recent example is based on continuous wave (CW) lasers that cause a phenomenon known as thermocavitation to create liquid jets by heating the injectate above its boiling point with an explosive phase transition.^8, 9, 27, 28, 35^

This work evaluates a CW-based needle-free micro-jet injector as a possible transdermal delivery alternative with minimal damage to the skin structure. For the purpose of this study, the topical solution delivery will be primarily associated with diffusion processes. In terms of delivery by the penetrating solid needle or liquid jet and subsequent diffusion, we will refer to this process as penetration. The combined diffusion and penetration processes are defined as total dispersion. We investigated the potential to achieve deeper dispersion depths than with topical application or solid needle injections. All three methods had different injection or application durations but were compared after 60 minutes for analysis. The metrics for comparison used in this work are the diffusion and dispersion depths, widths and surface areas for the topically applied or injected solutions, and the evaluation of the spread of the drug within the skin, characterized by the dispersion distances.

## 2. Materials and Methods

Three different types of experiments were performed with porcine skin: one topical and two injection experiments. For the first injection experiment, a permanent make-up (PMU) machine was used as solid needle injector. The other was a needle-free micro-jet injector based on thermocavitation. For all methods, excess solution was carefully removed from the skin with a Kimwipe paper tissue (Kimberly-Clark).

### 2.1. *Ex-vivo* Porcine Skin

*Ex-vivo* abdominal porcine skin was used for the experiments from the abattoir (Slagerij Nijboer, Enschede, the Netherlands). Since the skin properties vary from animal to animal, all experiments and, replicates were made with the exact same porcine skin sheet to ensure a fair comparison of results. Fresh porcine skin samples were stored at 4°C in Dulbecco’s Modified Eagle Medium (DMEM, Sigma-Aldrich). A surgical knife was used to remove excess fat tissue and to cut the skin samples into 3 × 3 *cm* sizes. The skin samples were carefully dried with a Kimwipe before each experiment to prevent solution running.

### 2.2. Liquid Solutions

Two different solutions were used in all experiments. The aqueous solution consisted of 0.15% wt rhodamine B (Sigma-Aldrich) and 0.25% wt direct red 81 (Sigma-Aldrich). While rhodamine B was used for its fluorescing properties, the direct red 81 was needed to maximize the laser energy absorption for the needle-free micro-jet injector. For the glycerol solution, 10% glycerol (Sigma-Aldrich) was added as diffusion enhancer due to its use in the pharmaceutical industry as a moisturizer.^15^ As measured elsewhere, both solutions behave as Newtonian fluids with measured constant viscosity η_aqueous_= 0.9 *mP as* and η_glyc10%_ = 1.2 *mP as*.^27^ The absorption was measured for both solutions in the spectral range of [300-1000] *nm* (UV-Visible Spectrophotometer UV-2600, Shimadzu).

### 2.3. Topical Application

A pipette (Eppendorf) was used to apply 2 *μl* solution drops onto the skin surface. Dye diffusion was compared at three different time intervals (after 5, 30 and 60 minutes). Once the time elapsed, any excess solution was carefully removed with a Kimwipe before being frozen in 2-Methyl Butane (Sigma-Aldrich). Each experiment was repeated five times, which resulted in 30 samples for the two different solutions.

### 2.4. Solid Needle Injector

The handpiece of the PMU machine (PL-1000 Mobil, Permanent Line GmbH) was vertically placed in a custom-made holder to inject various solutions into porcine skin, as shown in Figure 1. The encased motor in the handpiece smoothly moves the solid needle within the attached cartridge in a cyclic pattern up and down.^39^ With the electrical control unit of the PMU machine, the frequency of the needle movement is adjustable in the range of f = [50 − 150] *Hz*. The disposable needle cartridge contains a sterilized, stainless-steel needle with a diameter of d_Needle_= 0.4 *mm* (Permanent Line GmbH). For the injection procedure, a syringe was used to dispense 2 *μl* of the solution into the cartridge orifice. The needle was fully retracted and placed at 1 *mm* distance from the porcine skin samples, which were fixed by an in-house 3D printed case.

**Figure 1:**
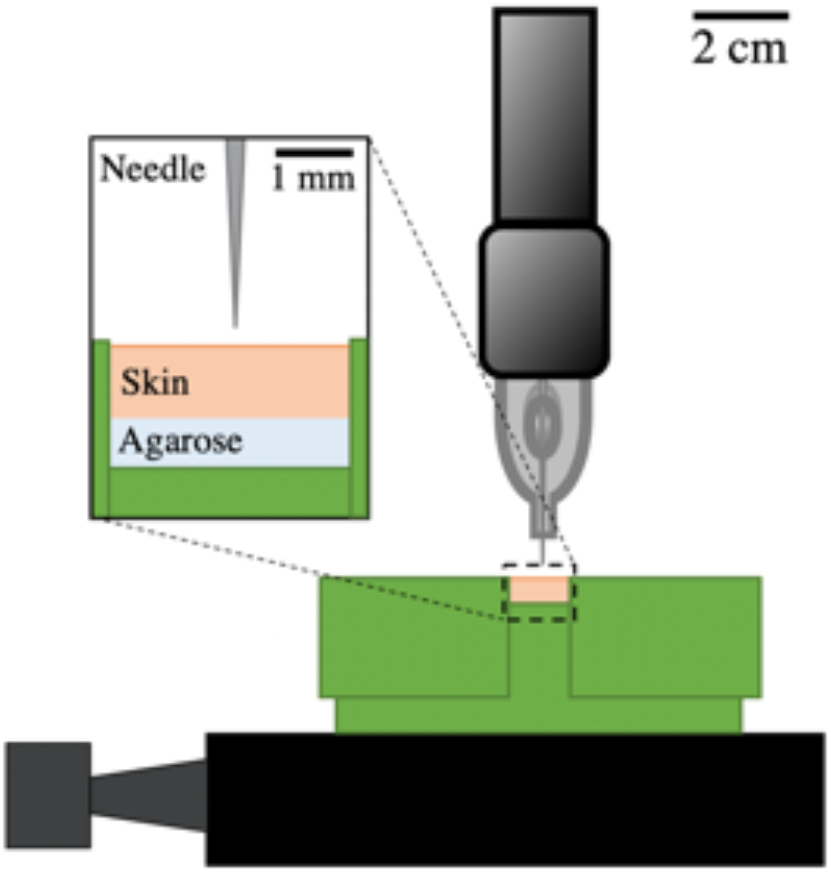
Schematic setup of solid needle injector. PMU machine hand-piece held vertically by custom-made holder for injections into porcine skin, which was fixed by 3D printed case.

To mimic the underlying muscle and fat tissue, 5 % wt agarose (Sigma-Aldrich) was prepared and poured into the 3D-printed holder. After the agarose solidified, the cut porcine skin samples were placed on top of it. Two lids held the skin tight, while the needle penetrated the skin at a frequency of 100 *Hz*. The injections of 20 seconds were timed with a circuit and an Arduino code, which turned on the PMU machine for the respective time. Five replicates were made for each solution, which in total resulted in ten porcine skin samples.

### 2.5. Needle-Free Micro-jet Injector

The experimental setup for the needle-free micro-jet injector consists of a CW laser diode (λ= 450 *nm*) and a microfluidic device composed of two anodically bonded borosilicate glass wafers (Schott AG). The fabrication process and design of these devices were previously described elsewhere.^8, 9^ The microfluidic devices consisted of two rectangular channels with 100 *μ*m depth: one for the cavitation bubble formation and the second one for the liquid jet to exit. An inlet of the same depth controlled the volume into the first channel (500 μm height, 1800 *μ*m length). The laser beam was focused with a 10x microscope objective at the bottom of the device, which was fixed with its holder to an XYZ linear translation stage holder to align the device with respect to the focused laser spot, as seen in Figure 2. Next, the first channel of the device was partly filled with one of the solutions by manually positioning the meniscus with a syringe to a channel position of 500 *μm*. A high-speed camera (Fastcam SA-X2, Photron) was then simultaneously triggered with a circuit and an Arduino code, which turned on the laser (U = 4.7 *V*, P = 1.2 *W*) for 100 *ms* to record all experiments at 160 × 10^3^ frames-per-second (fps). Within a few microseconds, the liquid inside the partially filled device was heated up above its boiling point. As a result, a fast-growing vapor bubble was created at the beam spot, a phenomenon known as thermocavitation.^28, 35^ The growing bubble simultaneously pushes the liquid out of the first channel and forms an injection jet. A white light source was positioned on the opposite of the high-speed camera to visualize the jet propagation (velocity and shape) and penetration into the skin samples. The jet velocities *v*_jet_ were calculated out of the image sequences of a single injection. Three experimental sets were made in which one sample either received one, three or six injections. Five replicates were made for each set and solution, resulting in a total of 30 samples.

**Figure 2:**
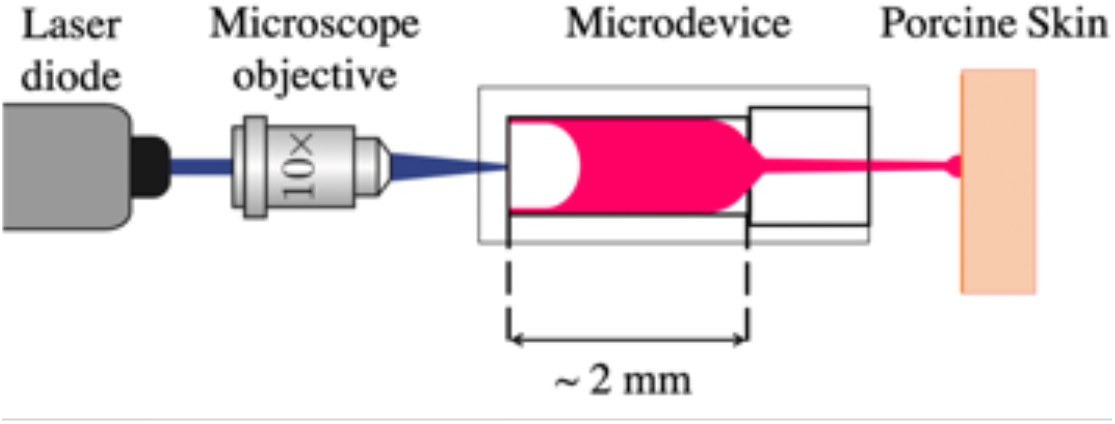
Schematic setup of the needle-free micro-jet injector. A microscope objective was used to focus the laser at the bottom of the microfluidic device. With a high-speed camera the bubble and jet formation, as well as skin penetration were recorded.

### 2.6. Analysis

After each topical or injection event, all samples were immediately embedded in an optimal cutting temperature mounting medium (OCT, VWR Chemicals) to stop the natural diffusion of the solutions in the skin. The samples were sectioned using a cryostat (Leica CM1950, Leica Biosystems) and analyzed under a Nikon E400 microscope for both bright field and fluorescence imaging. The solution diffusion throughout the different skin samples was quantified with the image processing software ImageJ, where the diffusion depth L_M_ and width L_T_ were measured 24 hours after the topical application or injection (see Figure 3). The applications of topical, solid or NFI themselves took less than 10 *s*, 10 *ms* or 0.5 *ms* respectively (Figure 3A). Further, the surface area of the injection site was determined with a MATLAB code developed for image processing. The fluorescence image was converted into a binary image, in which all the white pixels were added up to calculate the surface area of the injected solution (Figure 3B). Fluorescence images of the topically applied solutions were inverted with ImageJ and overlaid with the respective bright-field images.

**Figure 3:**
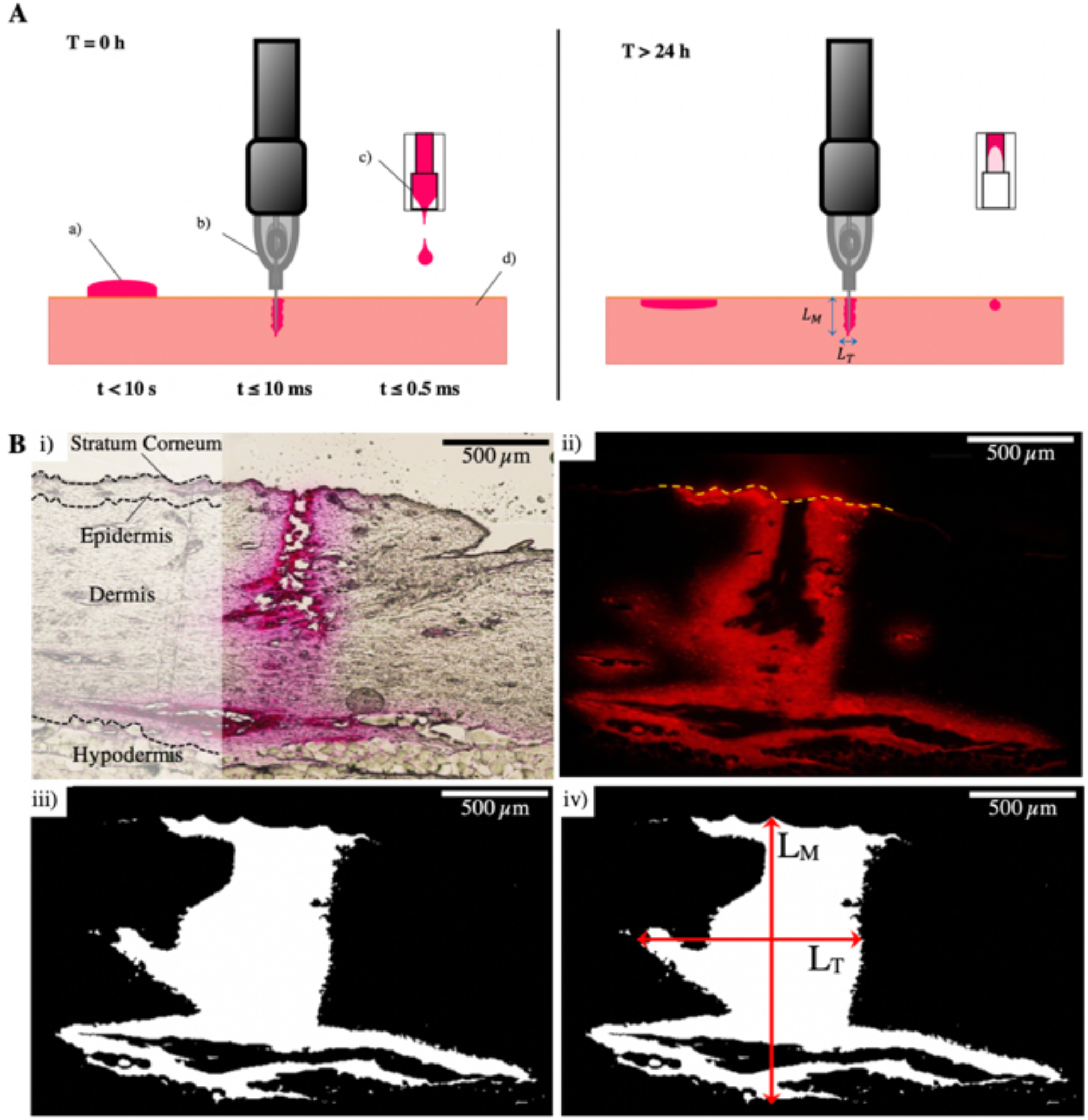
**A. Injection phases. Left panel: Instantaneous application or injection** takes less than 10 *s* for **a)** topical application and **b)** less than 10 *ms* for solid needle injection or **c)** 0.5 *ms* for micro-jet injection. **Right panel: Solution dispersion** throughout the skin, 24 h after application/injection. **d)** porcine skin with the stratum corneum; **B. Image processing analysis. i)** Bright-field image of a solid needle injection and schematic drawing of different skin layers showing the stratum corneum on top. Glycerol solution was injected for 20 seconds at 100 *Hz*. **ii)** Respective fluorescence image; the yellow dotted line indicates the porcine skin surface. **iii)** Calculated surface area of glycerol solution injection represented in white after image processing (101 × 10^4^ *μm*^2^). **iv)** Diffusion depth **L**_**M**_ and width **L**_**T**_ measurements with ImageJ. Scale bars correspond to 500 μm.

## 3. Results

### 3.1. Topical Application

The dye diffusion after the topical application was first analyzed. Rhodamine B, shown in Figure 4 in light blue, diffused 44-64% deeper into the skin than the non-fluorescing direct red 81 (pink), which remained more superficial. The direct red 81 diffusion of the glycerol solution, however, seems to be higher than in the aqueous one. Especially after 60 minutes, in which the non-fluorescing part of the glycerol solution roughly diffused twice as deep as within the aqueous solution. The average diffusion depth (in *mm*) of the fluorescing rhodamine B in both solutions was calculated from the respective samples for each time interval after 5, 30 and 60 minutes (see Figure 4). In all measurements, the topical application of the glycerol solution achieved 7-29% deeper diffusions than the aqueous one. However, there was no statistically significant difference between the two solutions. The average vertical diffusion rate calculated between the three-time intervals show that the glycerol (*dz*/*dt* = 0.07 *μm* /*s*) and aqueous solution (*dz*/*dt* = 0.05 *μm*/*s*) diffused deeper with time, while the lateral diffusion of both solutions was twice as big. The widest experimental scatter was seen with the glycerol solution after 60 minutes of topical application.

**Figure 4:**
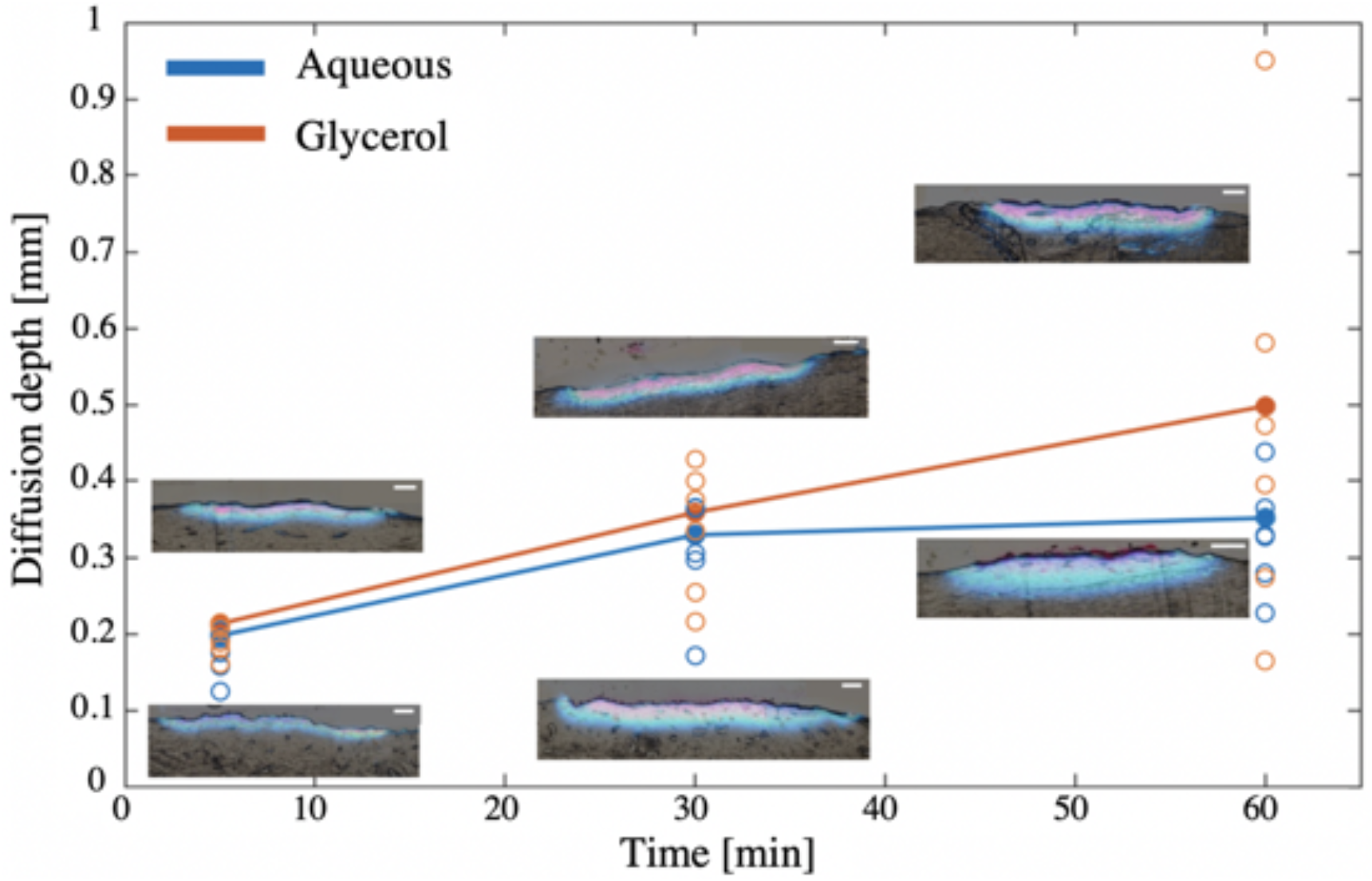
Diffusion after topical application. Average diffusion depth after 5, 30 and 60 *min* of topical aqueous (blue) or glycerol (orange) solution application as shown by filled data points. Other data points represent the experimental scatter of all 30 samples. Inverted fluorescence (blue) and enhanced bright-field (pink) image overlay of topical diffusion.

### 3.2. Solid Needle Injection

Under similar conditions as reported above, the dye dispersion of both solutions was studied after solid needle injections with the static mode. The surface area of the dispersed solutions was measured with a MATLAB code and is displayed by the surface areas of the circles, as seen in Figure 5. The averaged depth, width and area results are represented by the filled circles respectively. The dispersion depth of the glycerol solution was almost as deep as its dispersion width (1.13 ± 0.6 mm). The aqueous solution, on the other hand, remained more superficial with 1.04 ± 0.2 mm and instead, dispersed more in the horizontal direction (1.28 ± 0.3 mm). Two fluorescent images of the respective solutions show the dispersion throughout the skin. The injection path of the solid needle into the permanently damaged tissue can be seen by the dark area in the central injection region. On average, the aqueous solution achieved a surface area dispersion of 8.6 ± 3.2 × 10^3^ *mm*^2^, while the glycerol solution was slightly larger with 9.2 ± 4.8 × 10^−10^ *mm*^2^. The terminal vertical and lateral dispersion rates for the solid needle injection are, on the one hand, determined by the maximum depth or width of each injection and, on the other hand, by the solution dispersion in the tissue. With an injection time of 20 seconds, the vertical dispersion rate was *dz*/*dt* = 54 *μm*/*s*, while the lateral one was *d*y/*dt* = 60 *μm*/*s*. The corresponding injection video is shown in supplementary movies.

**Figure 5:**
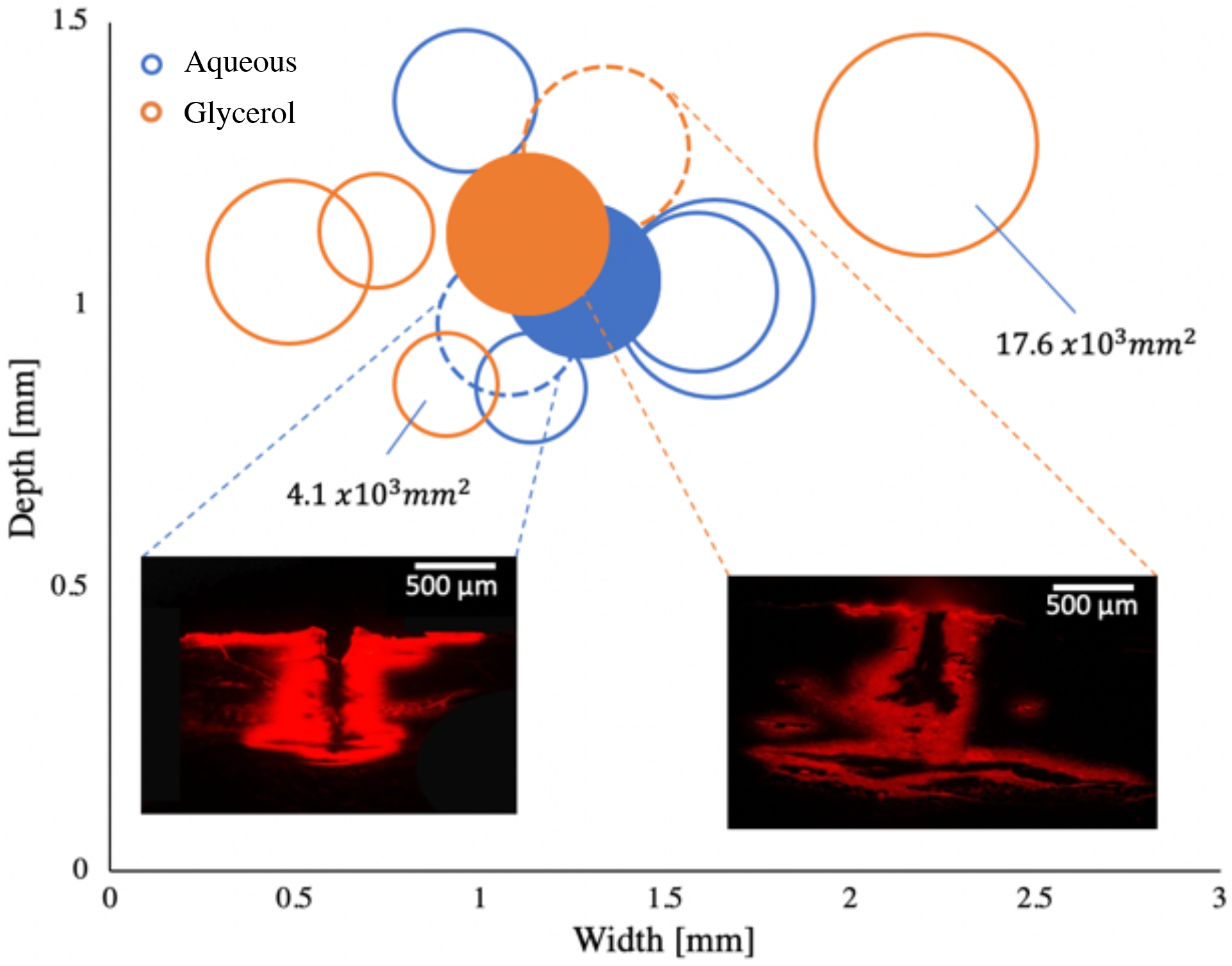
Solid needle injection with the static mode. Depth and width measurements of injections with aqueous (blue) and glycerol (orange) solution. Filled circles present the average measurements respectively. The circle sizes correspond to the measured surface area of the injections. Presentation of respective fluorescence images with the same exposure time of 150 *ms*.

We also performed more realistic dermal pigmentation injections in which the skin samples were set in motion along a plane perpendicular to the needle. More details of this experiment can be found in the supplementary information.

### 3.3. Needle-Free Micro-Jet Injection

The image sequence of an NFI, taken with 160 × 10^3^ fps, shows one out of three single injections that were made into one porcine skin sample (Figure 6A). The back-splash of the penetrating jet can be seen once it reached the skin (Figure 6A Bottom). Prior to the injections, the skin sample was placed at a 7.3 *mm* distance from the microfluidic device and the first channel was partly filled with 0.03 ± 0.003 *μl* aqueous solution. Once the laser is triggered, a vapor bubble was created inside this channel (Figure 6 Top). The aqueous solution cavitated within 75 ± 1.9 *μs* until the bubble pushes the liquid out of the first channel, while the glycerol solution needs more time (90 ± 1.3 *μs*) to form a jet of the same diameter (0.1 *mm*). With both solutions, however, the tip of the jets started to detach from the rest at 227 ± 18 *μs* of propagation. Between 329 ± 25 *μs*, all 5.5 ± 0.26 *mm* long jets completely left the microfluidic device. Directly after this, the jet started to break into droplets from the back of the tip in the direction of the jet displacement. The jet break-up of Newtonian liquids, such as water, are known to be caused by Rayleigh-Plateau instabilities.^14, 23^ At that time, the jet diameter decreased to 0.09 *mm* and remained the same until the first drop of the jet hit the porcine skin surface at 468 ± 80 *μs* of propagation. While the rest of the jet followed, 23.7 ± 10.4% of the solutions splashed back. Finally, the last drop of the injected solution reached the skin surface 274 ± 0.3 *μs* after the laser was triggered, while on average the injection process took 468 ± 80 *μs*. The corresponding NFI video is shown in supplementary movies.

**Figure 6:**
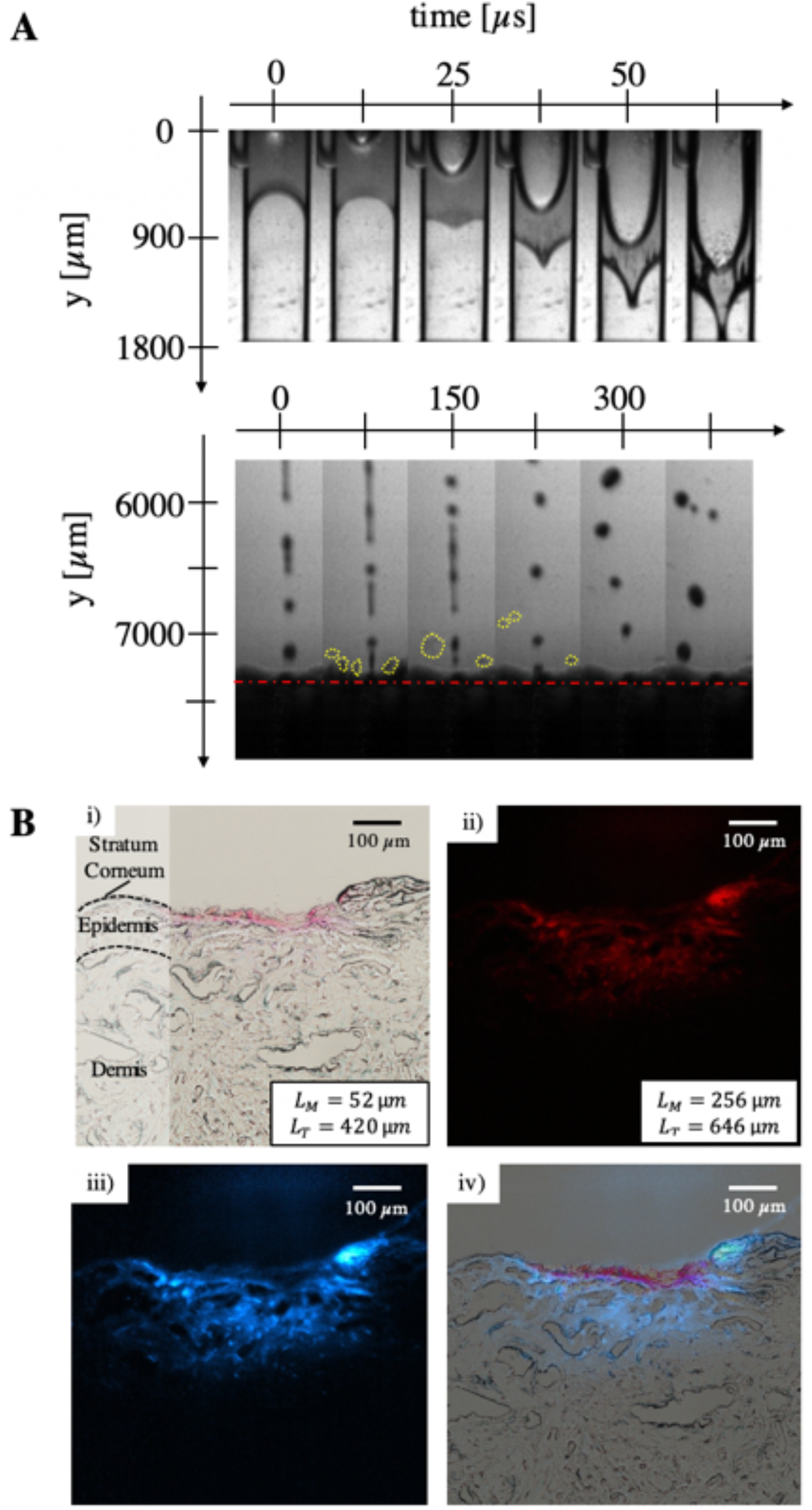
**A. Image sequences of NFI.** Recordings were made at 160 × 10^3^ *fps*. The jet velocity was 25 *m*/*s*. **Top)** Bubble formation and growth by thermocavitation within 62.5 *μs*. **Bottom)** Aqueous solution jet penetrates into porcine skin sample, which surface is indicated by the red dashed line. The yellow circles show the back-splashes of the jet. The distance between the microfluidic device and the skin sample is 7.3 *mm*. **B. Bright-field and fluorescence images of three repetitive, NFIs with the aqueous solution.** All scale bars equal to 100 *μm*. **i)** Bright-field image and schematic drawing of different skin layers. **ii)** Fluorescence image of solution dispersion. **iii)** Fluorescence image with ImageJ modified fluorescence color afterward. **iv)** Overlay of bright-field and fluorescence image.

The different filters and settings of the microscope used to analyze the injected tissue allowed to take bright-field, as well as fluorescence images of the solutions. The images of the specific sample shown above are presented in Figure 6B. The pink injection spot could be clearly seen in the bright-field image. Its size measurements are 0.45 *mm* in width and 0.09 *mm* in depth (Figure 6B i). The same sample was imaged with the fluorescence settings of the microscope. In comparison, the red area in Figure 6B ii) was almost three times as wide and twice as deep than the spot in the bright-field image, which becomes more evident in the overlay (Figure 6B iv). As for the fluorescence image of the solid needle injection, the fluorescence image had to be processed with ImageJ as well to have a higher contrast between the overlaying colors.

In the NFI experiment, the samples were differentiated by the type of injected solution and by the number of injections that were made into one sample. All data points in Figure 7A show the size and surface measurements for samples, which received one, three or six injections. The blue data points present the injection results of the aqueous solution, while the orange ones belong to the glycerol solution. Like the solid needle measurements in Figure 5, the circle sizes correspond to the measured surface areas of the dispersed solutions. The aqueous solution achieved greater lateral and vertical dispersions after one or three serial injections into one sample than the glycerol one. The differences of the dispersion depth, width and surface area between the solutions were less significant if six injections in a row were made into one sample. However, in all NFIs the glycerol solution resulted in a wider dispersion as in the topical application and solid needle experiments. Both terminal dispersion rates were determined as for the solid needle injection. The lateral dispersion rate (*d*y/*dt* = 20 × 10^5^ *μm*/*s*) was roughly seven times bigger than its vertical one (*dz*/*dt* = 3 × 10^5^ *μm*/*s*). Moreover, the velocities of all created jets are given in Figure 7B. On average, the glycerol solution jets were 2 *m*/*s* slower than the ones from the aqueous solution (*v* = 25 ± 2.3 *m*/*s*).

**Figure 7:**
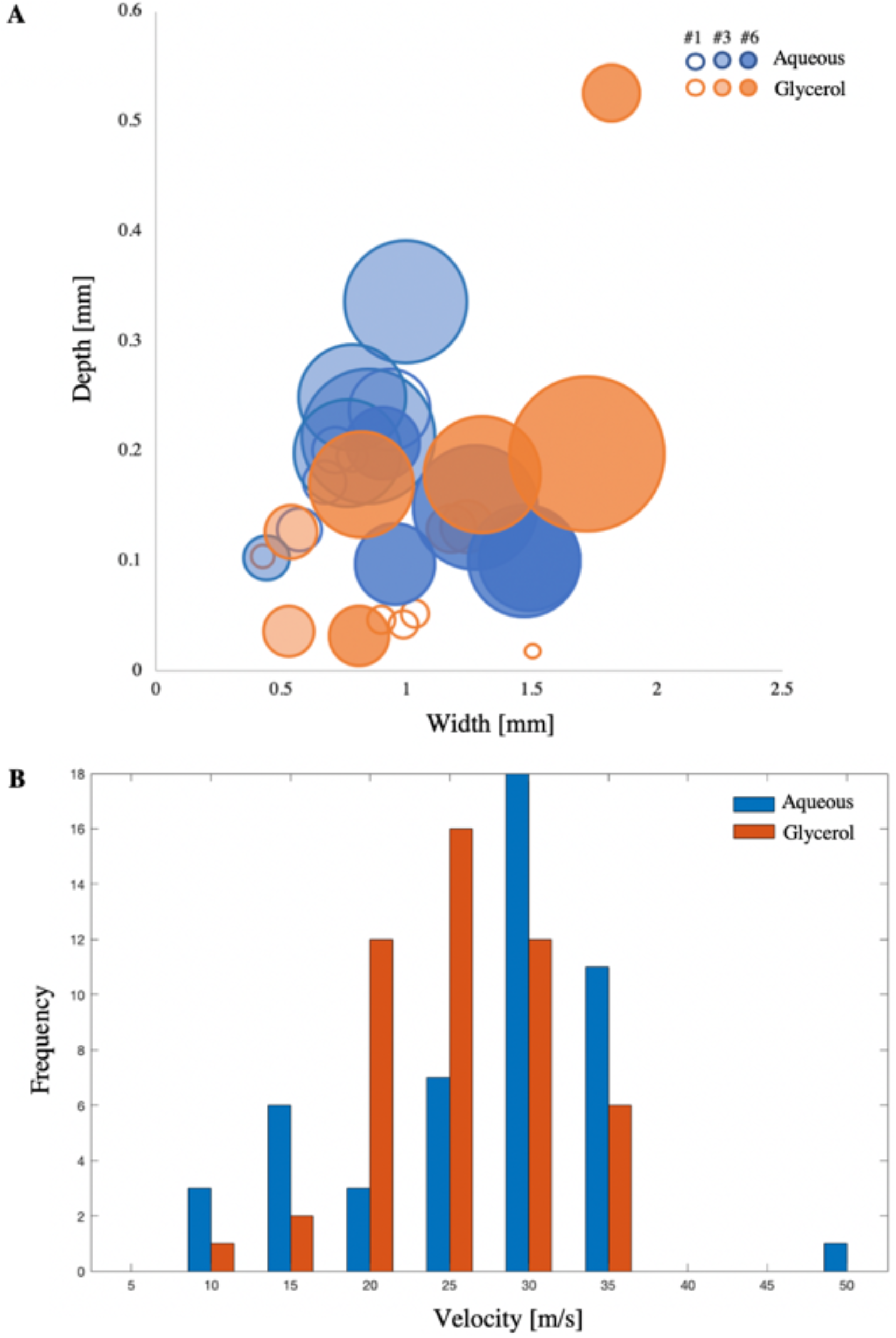
**A. Depth and width measurements of the needle-free injected solution penetration.** Penetration depth of aqueous (blue) or glycerol (orange) solution after one **(#1)**, three **(#3)** or six **(#6)** injections. Circle sizes correspond to the measured surface area of the injections. Biggest and smallest surface areas are indicated. **B. Velocity distribution of needle-free created jets.** The velocity occurrence of each aqueous (blue) and glycerol (orange) jet is plotted against the velocity.

## 4. Discussion

### 4.1. Aqueous and Glycerol Solutions

After the solution preparation, the coloration of both observed with the naked eye was identical. However, detailed analysis with a UV-Visible spectrophotometer revealed 49% higher absorbance with the aqueous solution throughout all measured wavelengths compared to the glycerol-containing solution. For the needle-free experiments, higher laser energy absorption led to faster heating of the aqueous solution, which resulted in faster bubble expansion and higher jet velocities. For the cases of single or three injections, the aqueous solution achieved deeper dispersion than that from glycerol, since higher jet velocities have more kinetic energy to penetrate the skin.

In all experiments and both solutions, rhodamine B exhibited wider and deeper diffusion throughout the skin compared to the non-fluorescing direct red 81. We attribute this difference to the diffusion coefficients of rhodamine B and direct red 81, but contributions for lower fluorescence detection limits cannot be ruled out. In general, the solution’s diffusion kinetics depend on two factors: the concentration gradient in the tissue and molecular weight. In this case, the latter factor plays the decisive role since molecules with high molecular weights (>500 *g*/*mol*), such as the direct red 81 (*M*_*W*_ = 675.60 *g*/*mol*), are known to have a poor transdermal delivery.^4, 10, 32^ The densely structured and outermost skin layer, the stratum corneum, additionally posed as a barrier for the direct red 81 macromolecules. While these were retained on the skin surface, the fluorescing rhodamine B molecules, possess a lower molecular weight (*M*_*W*_ = 479.02 *g*/*mol*) and thus, diffused more readily into the skin. The different diffusion kinetics became particularly evident if the solutions were topically applied and left on the skin for a longer diffusion time. The 7-29% deeper diffusion depths of the glycerol solution compared to the aqueous solution, however, are due to the different diffusion coefficients *D*.^11^

### 4.2. Comparison of Transdermal Delivery Methods

Topical application is the easiest and possibly the least invasive way to deliver molecules into the body. As mentioned before, the skin barrier limits this passive method to lipophilic molecules with a low molecular weight. Therefore, a diffusion depth of 0.35 *mm* or 0.5 *mm* is only achieved 60 minutes after the application of aqueous or glycerol solutions respectively. During the same time, 10 or 60 skin injections could have been made with the needle-free or solid needle method respectively. The calculated diffusion rate in vertical direction, *dz*/*dt* = 0.1 *μm*/*s*, was the slowest of all compared methods (Figure 8a).

**Figure 8:**
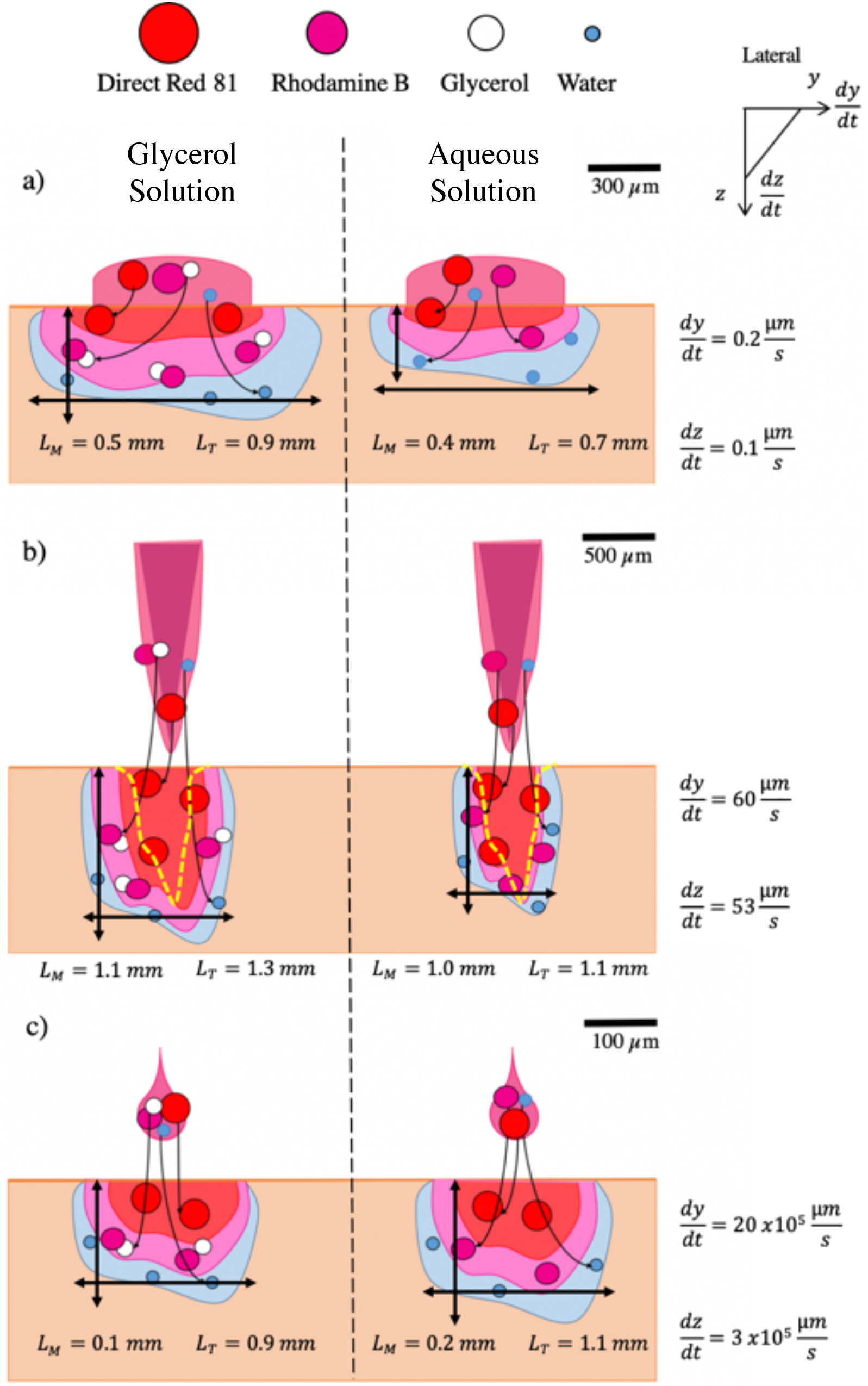
Schematic drawing of solution diffusion and dispersion in the topical application and injection methods. The left column shows the glycerol solution components diffuse or disperse differently across the skin, while the right one presents the aqueous solution. Average diffusion depth L_M_ and width L_T_ with the corresponding vertical, *dz*/*dt*, and lateral diffusion rate *d*y/*dt* for the respective methods. **a)** Topical application. **b)** Solid needle injection. The yellow dotted line indicates the skin damage caused by needle penetration. **c)** Needle-free micro-jet injection.

The solid needle injections with the PMU machine achieved the deepest penetration with a vertical penetration rate *dz*/*dt* = 54 *μm*/*s*. However, it was also the most invasive method. As seen in Figure 8b, needle penetration is detrimental to skin integrity, and is known to pose risks of infection and scarring.^4, 12^ The bright-field and fluorescence images show the ruptured tissue, which was caused by the needle penetration. However, this tissue damage did not always correspond to a deep dispersion depth of the solution. In terms of the experimental procedure, the adjustable PMU pen holder enabled a quick and repeatable injection process for multiple samples with the same penetration depth.

The injections with the NFI device are 20 times faster and lead to lateral and vertical diffusion rates larger than the solid injector, ca. 30.000 and 6.000-fold respectively (Figure 9c). The preparations for jet injections take more time since this setup is still experimental (30 minutes longer). Defects within the microfluidic devices, which appear during the production process, and gas bubbles that are formed in the channel if filled improperly, can scatter the incoming laser light. Less energy is absorbed by the liquid and as a consequence, the vapor bubble within the device does not grow rapidly enough to displace the liquid out of the channel. The image sequence in Figure 6Aii) shows that 6% (1.8 *nL*) of the solution jet splashed back from the skin surface. On average, 23.7 ± 10.4 % of the initial filling volumes did not penetrate into the skin due to splashbacks. According to the literature, a minimum jet velocity of 13 *m*/*s* should be sufficient to penetrate a typical skin strength of 20 *MPa*.^9, 29^ Out of all created jets, 92% were faster or even twice as fast than this velocity.

The most relevant observation is that no tissue damage was observed after analyzing the cross-section images. The stratum corneum remained intact even after six repetitive injections into the same sample spot. Since all excessive solutions were carefully removed from the skin after each injection and before freezing, we can assume that passive diffusion plays a negligible role. A comparison of the dispersion kinetics shows that the NFI method is three to six orders of magnitude higher than the solid needle and topical one respectively. In theory, the needle-free micro-jet injector is more efficient in terms of dispersion depth per second than the traditional methods. However, multiple injections with the needle-free device have shown that the solution dispersion does not increase linearly in time or number of injections. Further studies are required to investigate the actual dependency between a given number of injection events and the corresponding final dispersion patterns.

We calculated the kinetic energy K = 1/2mv^2^) of the needle and liquid jet that is transferred to the skin and evaluate the injection efficiency in terms of energy per injection *ε*_*energy*_ = *K*/*E x* 100 % (Table 1). The PMU machine has six and nine times higher *K* and *ε*_*energy*_ respectively than the NFI device because of the larger mass. The penetration strength S = p/E indicates potential skin damage by quantifying the skin rupture and penetration characteristics. This dimensionless variable is the strength ratio between the injection pressure p exerted onto the skin and tabulated skin’s Young’s modulus *E*_*Skin*_ = 0.5*MPa*.^7^ The jet pressure is calculated as 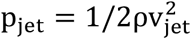 with *ρ* as the liquid density and *v*_*jet*_ as the jet velocity.^37^ The needle pressure value (*p* = 2400 *kPa*), taken from a similar experiment with agarose, is assumed to be within the same order of magnitude for porcine skin.^27^ Both solutions in the NFI experiments resulted in a nine times lower penetration strength than the solid needle (*S* = 4.8). Despite their low *p* and thus, *S* values (*S* < 1), which theoretically implies no penetration, the bright-field and fluorescence images confirm the jets entered the skin.^38^ We hypothesize that the Young’s modulus of these skin samples is lower than obtained from literature, since the moduli vary for different skin samples and are highly dependent on the anatomical skin location and measurement technique, such as stress-strain or indenting experiments.^2, 7, 13, 41^

**Table 1:**
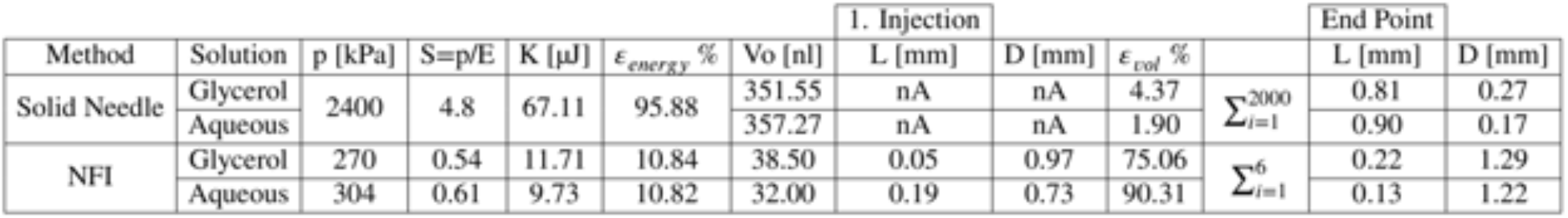
Comparison between the solid needle and needle-free micro-jet methods. The ratio between the injection pressure *p* and the skin’s Young’s modulus *E*_*Skin*_ = 0.5*MPa*,^7^ defines the penetration strength *S*. The diffusion depth *L*, width *D*, and volume *V* are quantified for a single NFI and for the endpoint. In the case of the solid needle, no data was available for one injection since the shortest injection time within these experiments were 20 s.

As for the efficiency of volume injected per injection event, the ejected (NFI) or deposited volume (solid needle) *V*_0_ is set in relation with the volume remaining in the skin as *ε*_*vol*_ = *V*_*inj*_ /*V*_0_ *x* 100 %. The following volume estimations are supposed to be taken as an indication, while future studies should focus on more precise measurements. The *ε*_*vol*_ of the NFI device was 19-45-fold higher than the PMU machine since the solid needle only injected 2-4% of its initial filling volume into the skin. The NFI device, on the other hand, injected 75-90% of its volume. The methods used to obtain these values are described in more detail elsewhere.^27^

In real-life conditions, repeatability or reproducibility of multiple injections may require that the jet injection frequency is increased to larger values than possible with the current experimental setup. For future *in-vivo* studies, we consider that the temperature of the liquid jets has to be measured to assess any probable burn injuries of the skin. To this aim, a thermographic high-speed camera could be used. As a next step, the caused pain needs to be evaluated in *in-vivo* studies, as it can be rated subjectively by each individual. Further, for future injections of medications, the molecular structure of the medication is not expected to change due to laser exposure and thermocavitation. The UV-visible spectrophotometer and proton nuclear magnetic resonance (^H^NMR) could be used to detect any structural changes and by-product generation due to heat.

## 5. Conclusion

The comparison of three transdermal delivery methods based on topical application, solid needle and, needle-free micro-jet injection, has shown that the latter was able to achieve a dispersion depth comparable to the topical application. The needle-free micro-jet injection has a faster injection and higher dispersion rate, which makes it a more efficient transdermal delivery method than the traditional ones. Moreover, no tissue damage was observed unlike the results with a solid needle injector. Since the needle-free micro-jet injector is still in an early development phase, experiments with human skin are needed to investigate if the created liquid jet is able to penetrate causing minimal damage. In theory, a needle-free injection might penetrate less deep into human than porcine skin due to the different and larger Young’s moduli of the stratum corneum.^21, 33, 34^ It is assumed that the penetration depth of the solid needle injection is not affected due to its high penetration strength S, which is defined by the ratio between injection pressure and Young’s modulus.

It can be concluded that the needle-free micro-jet injector has the potential to improve drug delivery across the skin if the challenges mentioned above are overcome. Millions of people that currently use painful solid needles for medical or cosmetic applications could soon replace them with a less invasive and safer liquid injection method.

## Acknowledgement

We would like to thank Dr. Loreto Oyarte-Galvez for her assistance with experiments, Stefan Schlautmann and Frans Segerink for their technical support with the microfluidic devices and the optical setup construction. The material support of PERMANENT-Line GmbH Co. KG. and MT-Derm is kindly acknowledged. DFR acknowledges the recognition from the Royal Dutch Society of Sciences (KHMW) that granted the Pieter Langerhuizen Lambertuszoon Fonds, 2016.

