## Supplementary for "A Comparison of Drug Delivery into Skin Using Topical Solutions, Needle Injections and Jet Injections"

**Supplementary Information**

1. Moving Mode

The skin samples were placed on a one-axis motorized translation stage. The time needed for the stage to cover a distance of 48.5 $cm$ was measured for each current. Ten measurements were made to estimate the average velocity. For each injection, 2 $\mu l$ of the solution was dispensed into the PMU cartridge. Along a plane perpendicular to the needle, the samples were set in motion (Figure 9). To ensure a straight injected line, the samples were fixed with two 3D-printed holders onto the stage. The PMU machine ran in the same frequency as in the static experiment and was manually turned on when the stage started to move from its starting point 15 $cm$ from the position of the PMU machine. The injections were made at four different velocities by varying the current I = [0.2, 0.3, 0.4, 0.5] $A$.

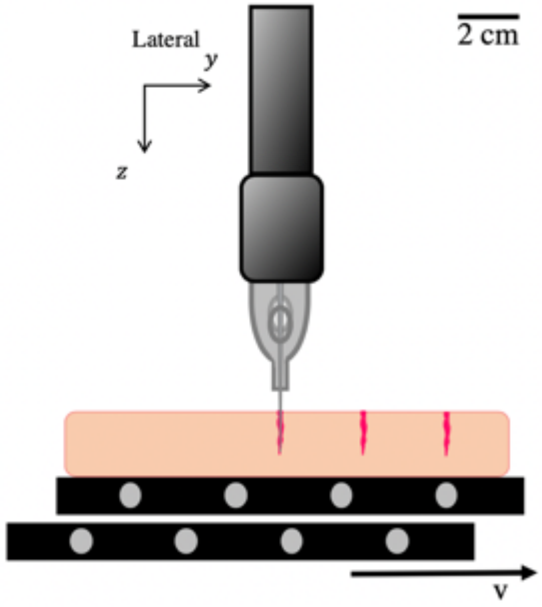


Figure 9: Moving injection. Porcine skin placed on a one-axis motorized translation stage with velocity $v$ ranging from [6 - 19] $cm/s$. PMU machine handpiece held vertically by a custom-made holder.

The same methods were used to quantify the dye dispersion after injection within the moving mode. The slowest velocity was measured for the lowest linear motor current of 0.2 $A$, corresponding to a displacement of 6 $cm/s$, while the fastest average velocity of 19 $cm/s$ was achieved with the largest current 0.5 $A$. The bright-field image in Figure 10A shows characteristic injection sites made with the moving mode. For these injections, the linear motor moved the stage and skin sample with 10 $cm/s$. The five injection sites are highlighted in the dashed boxes, while the numbers indicate the injection order. A typically measured distance of 0.85 *mm* between two injection sites is pointed by the arrow.

Injection results of the aqueous solution into the moving samples are shown in Figure 10B. Each graph is a result of an injection process into one skin sample that was moved at a velocity according to the current of the linear motor. Since each data point shows one injection site, the distances between them represent the distances between the respective injection sites. All dispersion depths were measured from the fluorescent images because the injection sites in the bright-field images were not visible for 0.4 *A* and 0.5 *A*. The dashed yellow graph shows measurements that were taken from the bright-field image shown above. A direct comparison between the two 0.3 *A* measurements shows that a deep needle penetration in the bright-field image does not necessarily correspond to a deep fluorescence dispersion or vice versa. On average these five injection sites were around 0.8 *mm* apart from each other. In contrast, larger distances of 1.2 *mm* were measured on the skin sample that moved faster at 19 $cm/s$. Due to the higher velocity, only three injections were made into the equally sized sample. No distinguishable single injections could be made into skin samples at 6 $cm/s$ because of a too slow displacement resulting in one injection site. The corresponding video of moving injection is shown in supplementary movies.

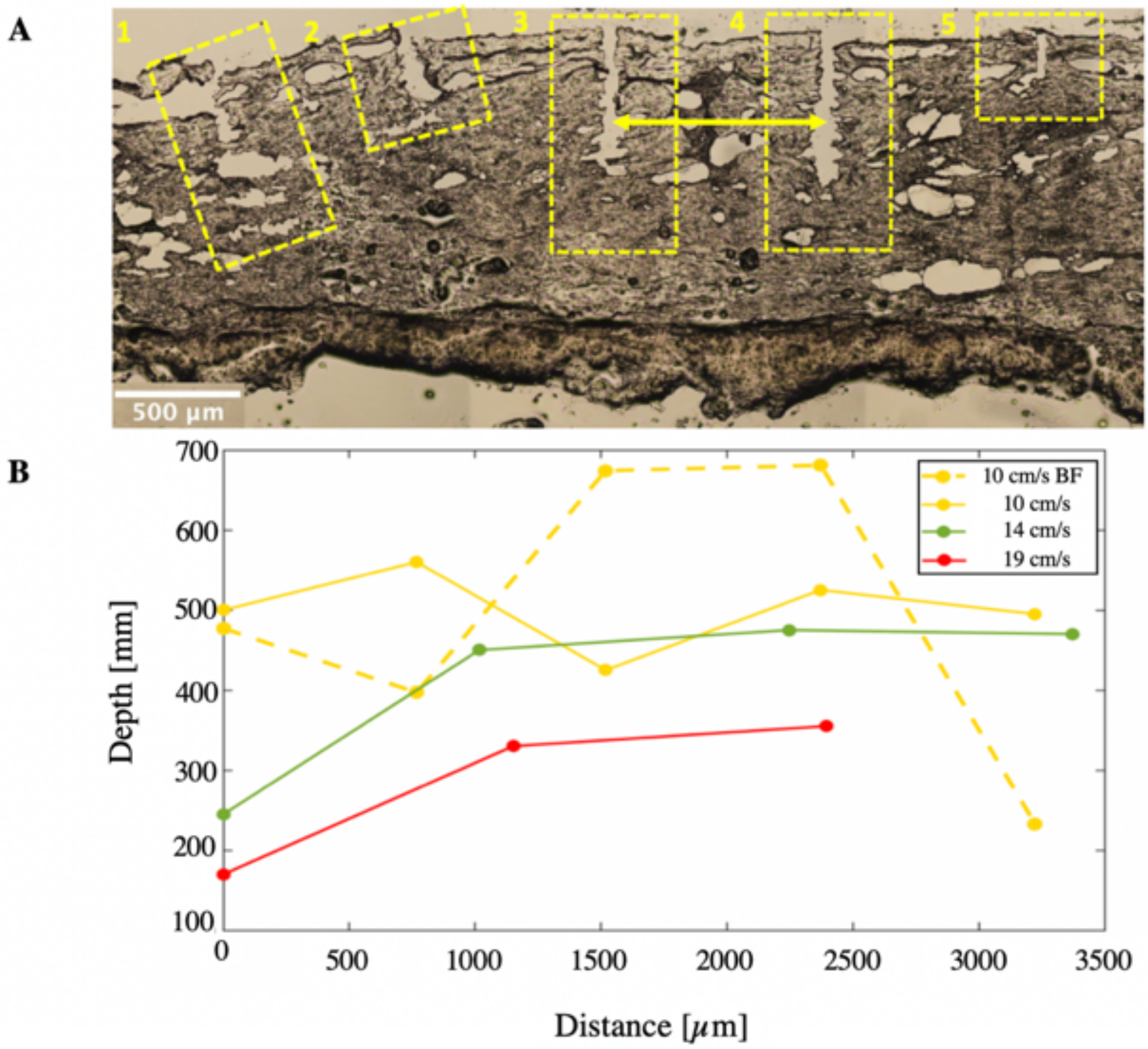


Figure 10: **Moving Mode. A.** Bright-field image of injections made into moving skin samples. The linear motor moved the stage and skin sample with 10 $cm/s$. The dashed boxes show the single injection sites made by the penetrating needle, while the numbers indicate the injection order. The yellow arrow shows a distance measurement between the injection sites. **B. Depth and distance measurements**. Skin samples were moved at different velocities $v$ = [10.0, 14.0, 19.0] $cm/s$ (yellow, green, red). The dashed yellow graph shows depth measurements made within the bright-field image in A. Depth measurements in fluorescence images are shown by continuous graphs. The dispersion depth is plotted over the distance between the injection sites, which are represented by the data points.

2. Continuous Wave Laser Diode Calibration

A power meter was used to measure the power of the laser diode. Therefore, the entire laser spot had to be captured by the detector of the power meter. An Arduino code turned the laser on for 30 seconds. Starting with 5.5 *V*, the voltage of the laser was decreased for each measurement by 0.1 *V* until a power of 0 *W* was measured.

The power of the laser was measured as a function of the voltage. The measurements started at 5.5 V (1.755 W) and reached 0.005 *W* at 3.5 *V*. The laser power has a linear dependency on the used voltage (Figure 11), which is shown by the blue data points approximating the fitted linear equation. The red dashed line indicates the used voltage of 4.7 *V* and the respective laser power 1.12 *W*.

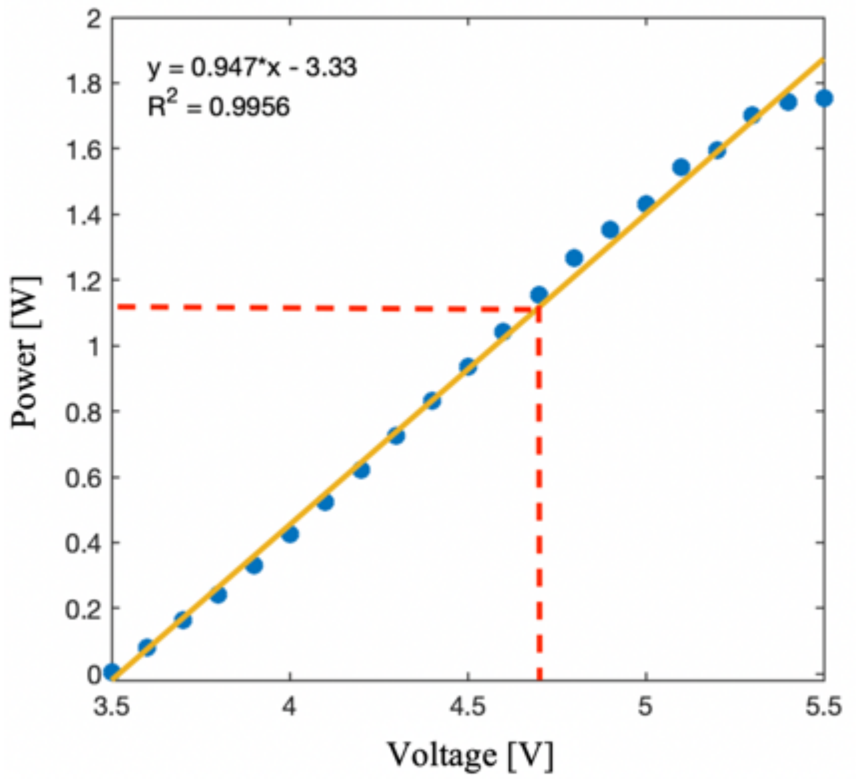


Figure 11: **Measured laser power (W) versus voltage (V).** Fitted linear equation (yellow) with regression $R^2$ = 0.9956. Used voltage 4.7 *V* corresponds to 1.12 *W* laser power.
