## Supplementary Videos for "A Comparison of Drug Delivery into Skin Using Topical Solutions, Needle Injections and Jet Injections"

**01_PMU_Form.W.**

The fully retracted needle was placed at 1 𝑚𝑚 distance from the porcine skin sample, which was fixed by an in-house 3D printed case. Injections of 20 seconds were timed with a circuit and an Arduino code, which turned on the PMU machine for the respective time. The recording was taken with 160 × 103 fps.

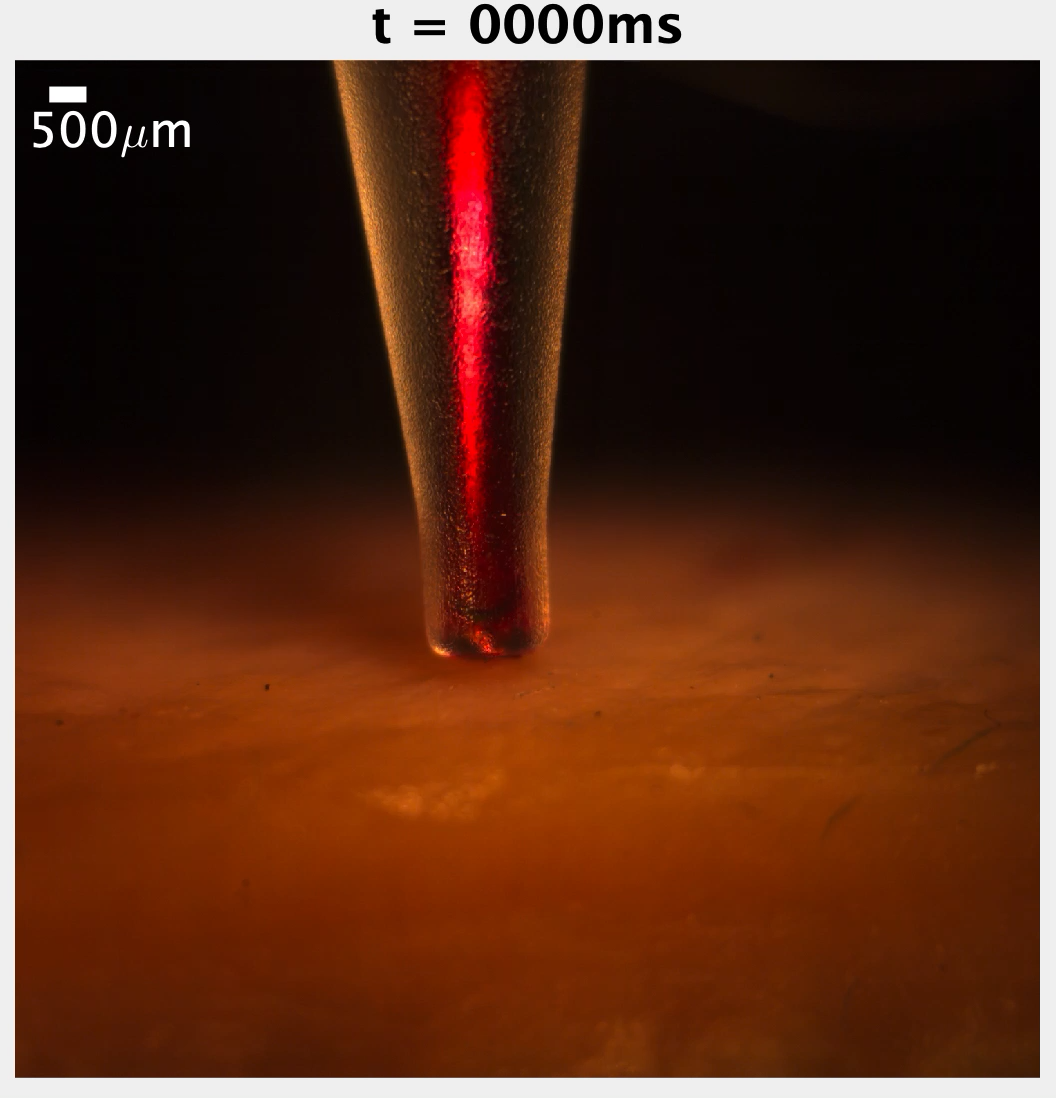

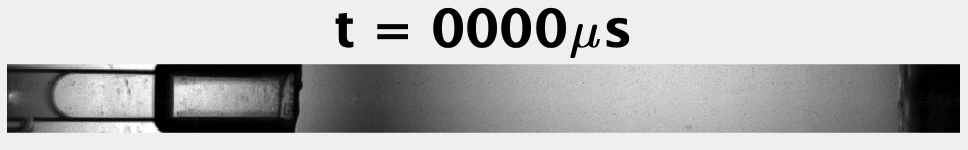

**02_NFI_Form.W.**

The recording of an NFI, taken with 160 × 103 fps, shows one out of three single injections that were made with the aqueous solution into one porcine skin sample. The bubble formation and growth by thermocavitation was within 62.5 𝜇𝑠 and resulted in a jet velocity of 25 m/s. The back-splash of the penetrating jet can be seen once it reached the skin, which was placed 7.3 mm from the microfluidic device.

**03_Dynamic_Form.W.**

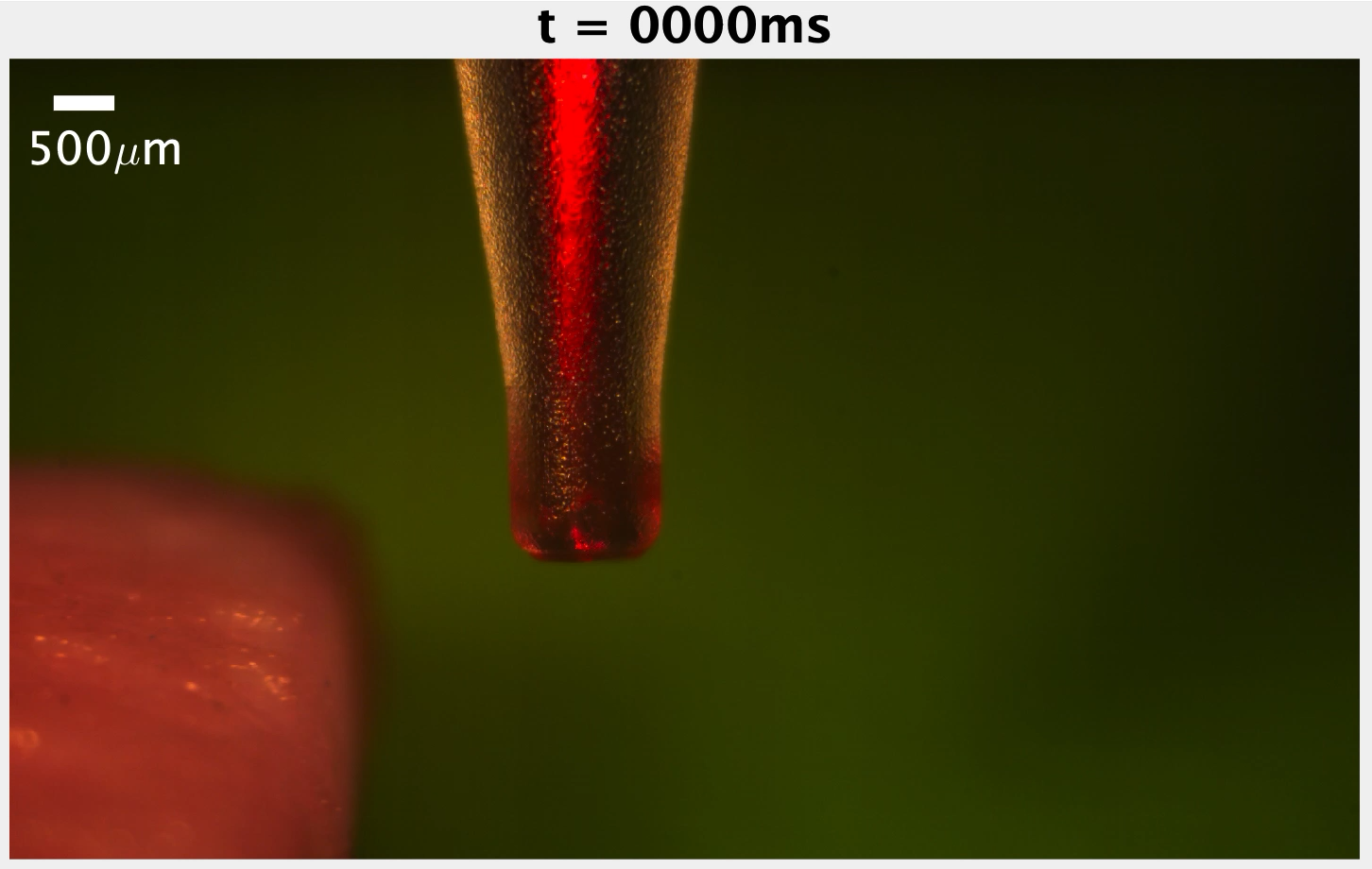
Along a plane perpendicular to the needle, the porcine skin samples were placed on a one-axis motorized translation stage and set in motion (v=10 cm/s). The PMU machine injected the aqueous solution at a frequency of 100 Hz. The recording was taken with 160 × 103 fps.
